# MINDFUL: A Method to Identify Novel and Diverse Signals with Fast, Unsupervised Learning

**DOI:** 10.1101/805820

**Authors:** Mallika Parulekar, Leelavati Narlikar

## Abstract

With rapid advances in experimental methods that map transcription start sites (TSSs) at a high resolution, there is a need to characterize the sequence diversity of TSS neighborhoods. Most current techniques scan for previously discovered elements, such as the TATA box, the INR motif, CpG islands, etc. to categorize promoters into different classes. Reliance on such elements hinders the discovery of novel elements. On the other hand, methods that use standard motif discovery to discover de novo promoter elements are also limited by the fact that a motif is picked up only if it is over-represented in the dataset. An element that appears only in a small set of promoters can thus be missed. We previously developed a clustering-based approach that uses no prior knowledge of elements to solve this problem [1]. That method uses Gibbs sampling to learn the model parameters, but is untenable on large datasets. Here we propose a new, fast method called MINDFUL, that uses a greedy *k*-means-like approach to cluster promoters aligned by TSSs into diverse classes, while also learning the optimal value of *k*. It is general enough to be used for any data that has categorical variables, and is not restricted to DNA.

## Methods

We model this as a clustering problem. The input is a set of *n* DNA sequences *X*_*1*_, …, *X*_*n*_, pre-aligned based on their respective TSS’s. Each *X*_*i*_ has length *L*. We assume that each *X*_*i*_ belongs to one of *k* clusters (classes), which is denoted with a cluster label *y*_*i*_. The clusters are modeled as a probability weight matrix *M*, or a product of *L* categorical distributions over *A*, C, G, T. Specifically, the probabilities for cluster *r* are denoted by *M* _*r*_^*j*^, where 1 ≤ *j* ≤ *L*. These distributions as well as the *n* cluster labels are unknown.

We have adapted the standard *k*-means clustering to learn *M* using a two-step iterative algorithm in MINDFUL. Let *X*_*i*_^*j*^ be the base in position *j* in sequence *X*_*i*_, and let *size*_*r*_ be the current size of cluster *r*. The probability *M*_*r*_^*j*^ can be computed using the frequencies of each base in position *j* for all sequences currently assigned to cluster *r*. Then, the probability of a sequence *X*_*i*_ occurring given a model is defined as:

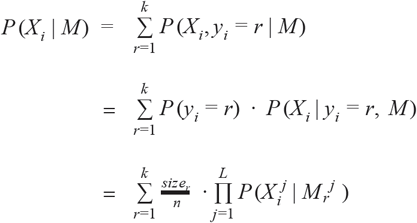

The pseudocode for finding the optimal cluster size is given below. We use 10-fold cross-validation and repeat the whole procedure with different initial points, to avoid local optima.

### Procedure *findKOptimal*

Repeat for *k=1* to *K*:

Let *H = {X*_*1*_,…,*X*_|*H*|_} be the 10% holdout and let *T* be the 90% training set Assign random clusters to sequences in *T* from *1* to *k* Repeat until convergence:

Recompute model *M* based on sequences in each cluster For each *X*_*i*_ in *T:*

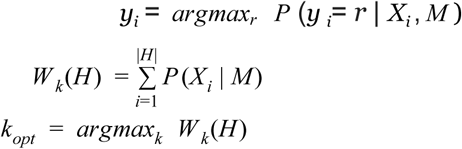

We finally set *k* = *k*_*opt*_ and repeat the clustering step until convergence for full set of sequences *X*_*1*_, …, *X*_*n*_.

### Binary Search Optimization

Instead of iterating over all *K* cluster candidates as done in *findKOptimal*, we use binary search, similar to previous work [2]. In each iteration we reduce the search space for *k*_*opt*_ by half, depending on *W*_*k*_ (*H*) values.

## Results

### Synthetic Datasets

We tested MINDFUL on several synthetic datasets, generated using various Dirichlet distributions. The probabilities of the four nucleotide bases in each position within each cluster were generated using the Dirichlet parameter **α**. We used varying **α** values to simulate noise; a higher alpha translates to more similar distributions of nucleotide bases across clusters, making it harder to compute the original clusters. The accuracy was assessed with the adjusted Rand Index (ARI), which was consistently in the range of 0.95 to 1.0 (with 1.0 being the highest).

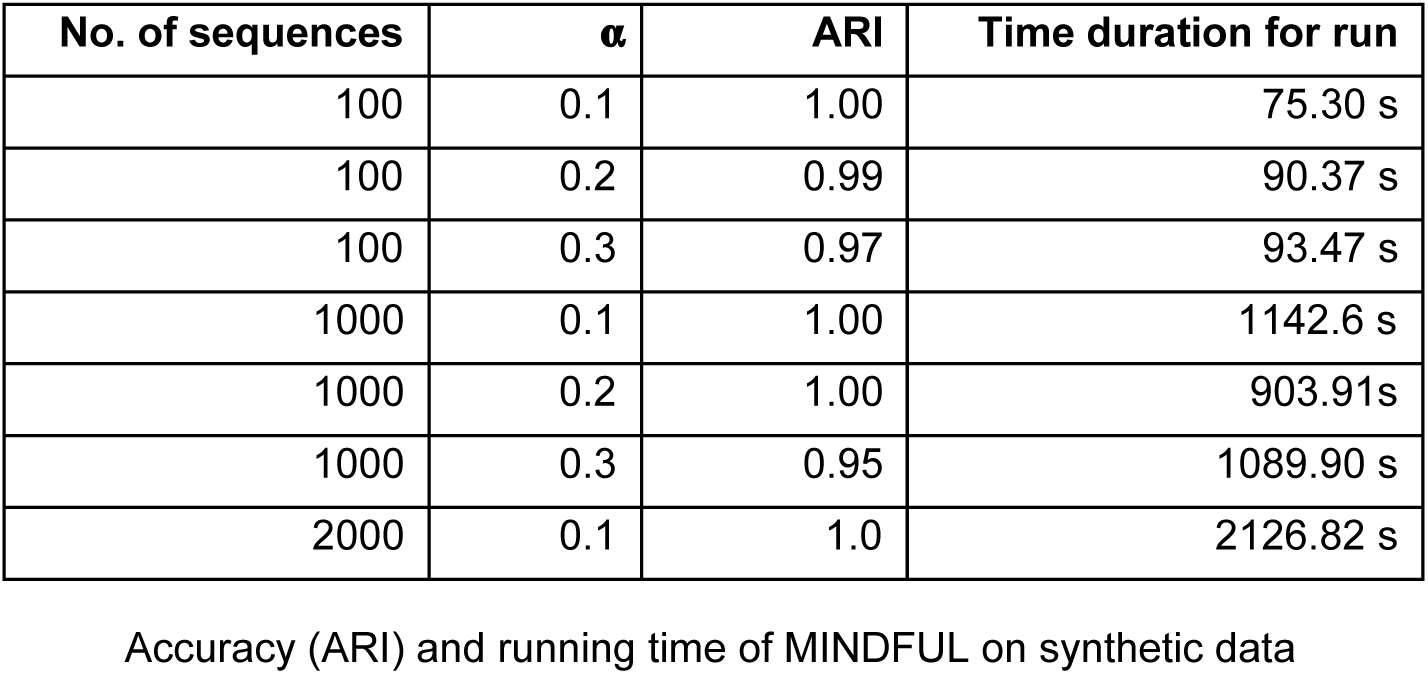

| No. of sequences | $\alpha$ | ARI | Time duration for run |
| --- | --- | --- | --- |
| 100 | 0.1 | 1.00 | 75.30 s |
| 100 | 0.2 | 0.99 | 90.37 s |
| 100 | 0.3 | 0.97 | 93.47 s |
| 1000 | 0.1 | 1.00 | 1142.6 s |
| 1000 | 0.2 | 1.00 | 903.91s |
| 1000 | 0.3 | 0.95 | 1089.90 s |
| 2000 | 0.1 | 1.0 | 2126.82 s |

Accuracy (ARI) and running time of MINDFUL on synthetic data

The figure below shows the clustering results for one synthetic data point.

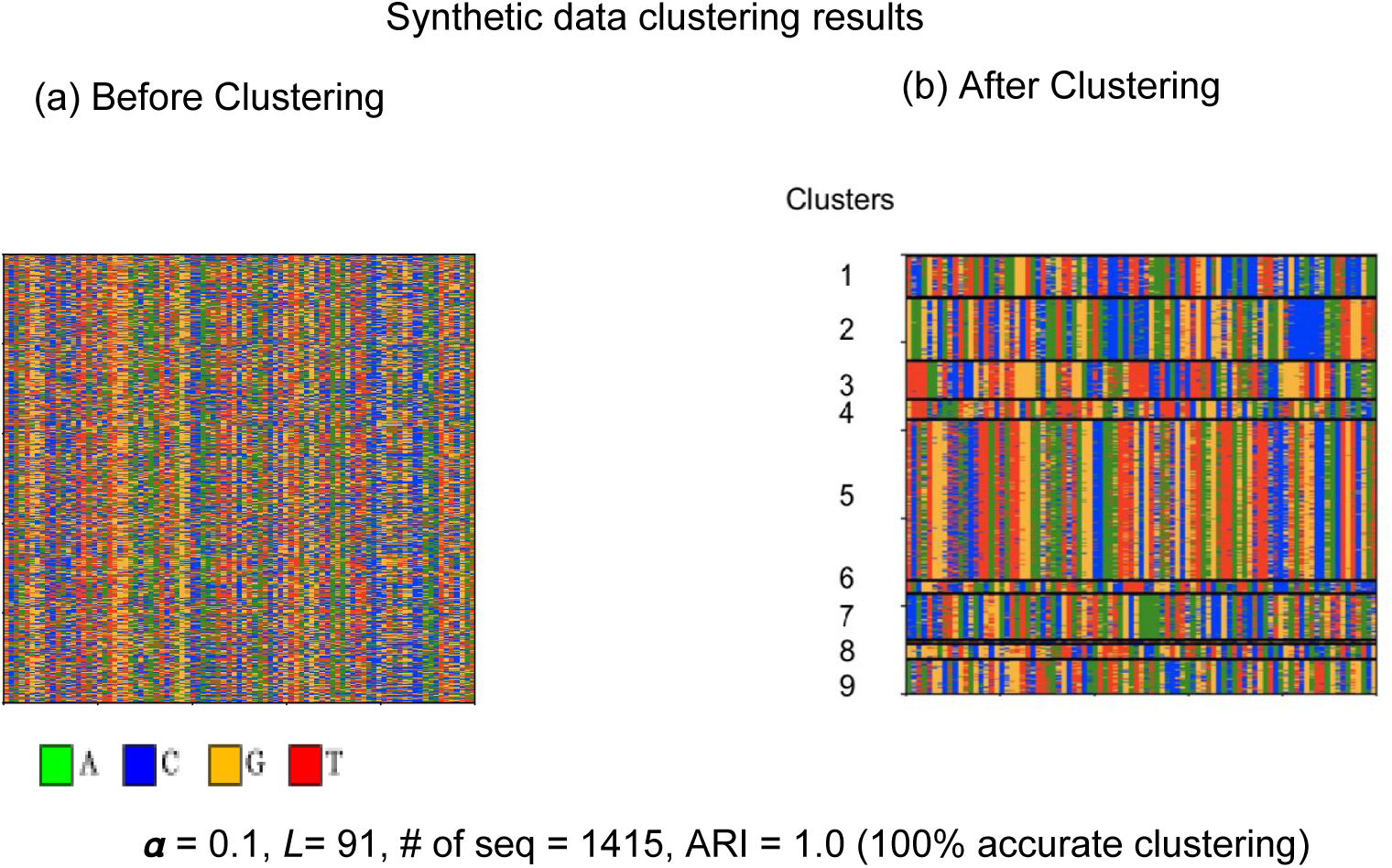

### Real Datasets

Next, we applied MINDFUL to fly [3] and tuberculosis [4] promoters. We were able to replicate past results of clustering on both these datasets. Our algorithm on the fly dataset achieved a speedup of *14.23x* over the best existing algorithm [2], which also uses unsupervised learning with binary search (see table below).

| Algorithm | Running time (Minutes) | Speedup |
| --- | --- | --- |
| Best existing ML algorithm | 401 | 1 |
| MINDFUL (no Binary Search) | 78.78 | 5.09 |
| MINDFUL (Binary Search) | 28.11 | **14.27** |

The clusters found by MINDFUL in biological data have distinct functional characteristics. For example, on the fly data set [3] shown below, we found that sequences from one of the clusters (cluster 2) were transcribed at a much higher rate than others. This cluster has the characteristic TCT motif at the TSS associated with ribosomal genes [5]. In the TB data, like the slower method, MINDFUL also identifies the TANNNT −10 box at varying distances from the TSS; each displaying a specific transcriptional profile [1].

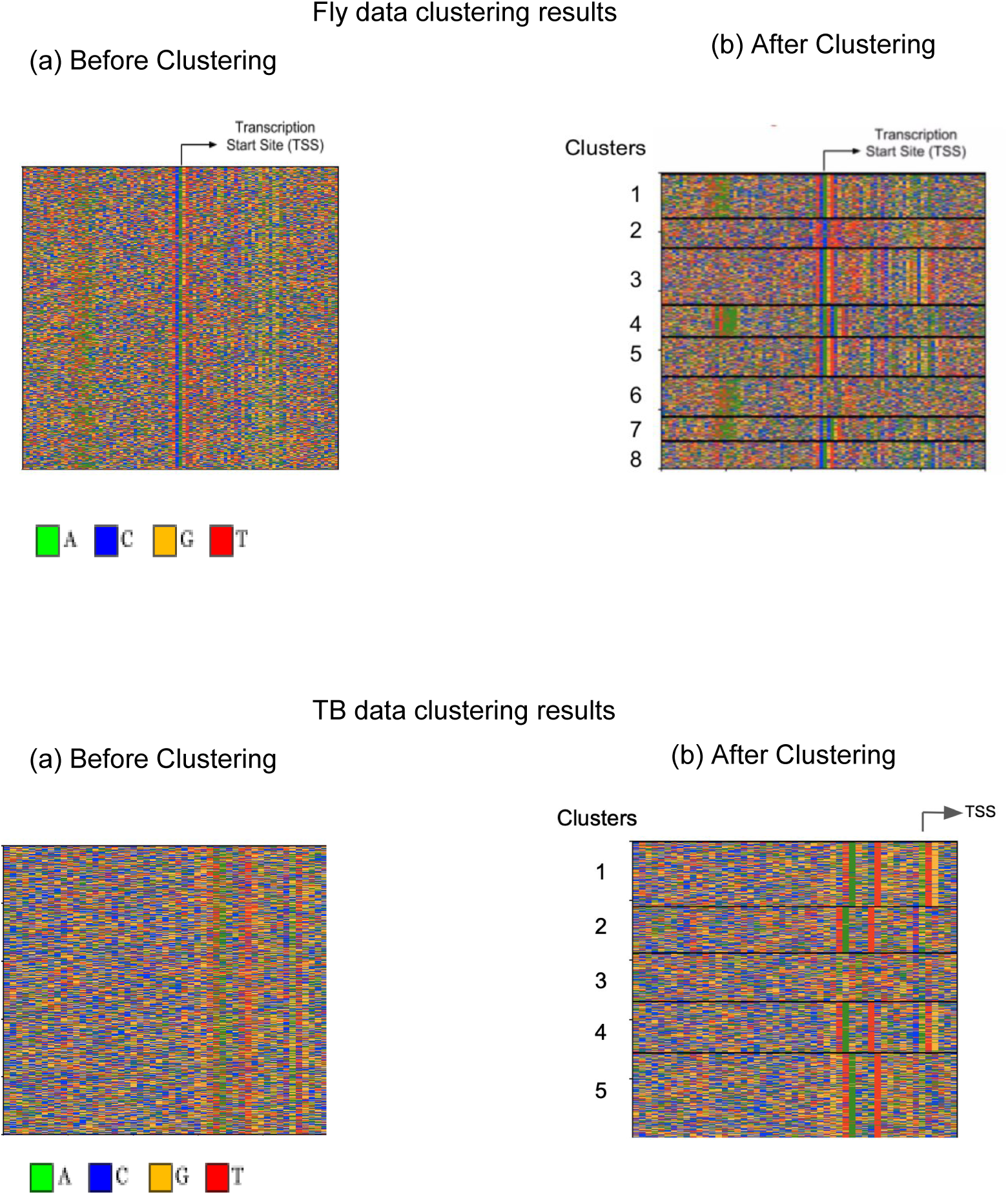

